# The C3HeB/FeJ Kramnik mouse as a model to investigate the efficacy of therapeutic tuberculosis interventions

**DOI:** 10.1101/2025.09.19.677270

**Authors:** Markus Depfenhart, Drieke van der Merwe, Suraj Parihar, Kobus Venter, Theunis Cloete, Yolandy Lemmer, Anne Grobler

## Abstract

Tuberculosis (TB) remains a formidable global health challenge, classified as the leading infectious disease killer, driven by the increasing prevalence of multidrug-resistant *Mycobacterium tuberculosis* strains. This persistent threat not only overwhelms public health infrastructures but also deepens socio-economic inequalities worldwide, underscoring the urgent need for novel therapeutic strategies. The C3HeB/FeJ Kramnik mouse model, frequently used for its capacity to mimic human-like granulomatous lesions, has emerged as a pivotal tool in TB research. However, despite its advantages, the model has limitations, particularly in terms of macrophage pathology and restricted B-cell activity, which pose significant hurdles for translating preclinical findings into human applications.

By developing models that better reflect human immune responses, the translation of findings into clinical applications may be improved, ultimately leading to more effective TB treatments and vaccines. This study aims to evaluate the Kramnik model by investigating the efficacy of a novel Pheroid^®^ formulation containing traditional TB therapeutics in the C3HeB/FeJ Kramnik mouse model. The potential and limitations inherent to the Kramnik model are explored and the need for complementary genomic analyses, such as sequencing SP110 and SP140, to elucidate the susceptibility linked to the SST1 locus is illustrated.

In addition, this article attempts to serve as a critical evaluation of the inherent limitations of current animal models like the Kramnik model, with the aim of paving the way for more accurate and reliable translational research in the fight against TB.

## Introduction

### Overview of Tuberculosis and its global impact

Tuberculosis (TB) remains a critical global health challenge, affecting millions each year, with the growing incidence of multidrug-resistant (MDR) and extensively drug-resistant (XDR) strains exacerbating the problem [1,2]. The World Health Organization estimates that approximately one-quarter of the global population carries latent TB, posing ongoing risks of reactivation and transmission [3], with associated socio-economic and humanitarian impacts, particularly in high-burden regions. In this complex landscape, reliable preclinical models are essential for advancing TB research and therapeutic development.

### Current treatment challenges and the development of resistance

The current treatment regimens for TB depend on drug sensitivity of the infective mycobacterial strain. The drugs used are classified into first- and second-line treatment regimens, based on efficacy, safety and side-effect profiles. Treatment is designed to reduce the development of resistance; it consists of a combination of several antibiotics taken over a lengthy period and is associated with numerous side effects that lower patient compliance [4]. This lack of adherence has been linked to the reinfection of patients and the emergence of resistant TB strains [5]. Furthermore, the current TB therapies are often limited by slow onset of action, extensive first-pass metabolism, and poor absorption of active pharmaceutical ingredients (APIs) [6,7].

Anti-TB drugs serve three main functions: they have bactericidal or bacteriostatic activity, their sterilizing activity is meant to prevent disease spread, and their use should prevent resistance. The first-line antimicrobials used for drug-sensitive TB are isoniazid (INH), rifampicin (RIF), ethambutol (EMB), and pyrazinamide (PZA) [8]. Among these, INH and RIF are the most potent bactericidal agents, effective against all populations of TB bacilli. INH inhibits the synthesis of mycolic acids, a crucial component of the bacterial cell wall, and is effective against actively growing TB organisms both inside and outside of cells [9,10]. RIF is a powerful sterilizing agent that targets slow-growing or non-replicating bacteria by blocking DNA-dependent RNA polymerase, thereby preventing mRNA and protein synthesis. PZA is thought to exhibit bacteriostatic or bactericidal effects depending on its concentration at the site of infection [9,10]. EMB, primarily bacteriostatic, plays a critical role in preventing resistance by blocking cell wall arabinan polymerization and interfering with multiple pathways such as RNA metabolism and phospholipid synthesis [9,10].

When first-line regimens fail and resistance develops, or when resistance to first-line drugs is detected, complex groups of second-line drugs are used, including fluoroquinolones (e.g., bedaquiline, ofloxacin, levofloxacin, moxifloxacin, ciprofloxacin), injectable agents like kanamycin, amikacin, and capreomycin, used in combination with EMB and PZA [11,12], the thioamides (ethionamide and protionamide), cycloserine plus terizidone or bedaquiline, delamanid and pretomanid [4,12]. These regimens are often hindered by high toxicity, cost, and the need for supervised administration to ensure compliance. Resistance to some of these drugs has also been reported [12], resulting in nonadherence. XDR-TB poses an even greater challenge, characterized by resistance to INH, RIF, any fluoroquinolone, and at least one second-line injectable drug [5]. Simplified, potent regimens that reduce therapy duration and minimize relapse are critically needed.

### The C3HeB/FeJ Kramnik mouse model and its significance in TB research

The lack of progress in the discovery and development of new anti-TB drugs over the past three decades can be ascribed to various factors, such as the business case for TB treatment development, compatibility issues between RIF and INH, and the complexities of shared drug toxicities and pharmacokinetic interactions in the case of Human immunodeficiency virus (HIV) co-infected patients. Furthermore, the lack of animal models available for testing drugs against latent and persistent *Mycobacterium tuberculosis (M.tb)* forms, and the absence of reliable biomarkers for evaluating new drugs are certainly contributing factors [13–15].

The C3HeB/FeJ Kramnik mouse model has become a standard in TB research due to its ability to replicate key aspects of human disease pathology [1,3,16–18]. The model offers a valuable platform for studying the complex interactions between *M.tb,* antimicrobials and the host immune system. The C3HeB/FeJ mice, named after Igor Kramnik, whose team developed the I/St mouse strain, so-named with reference to a specific genetic locus (Super-Susceptibility to Tuberculosis 1; SST1), have been found to possess a heightened susceptibility to *M.tb* infection with associated formation of well-defined hypoxic, caseous necrotic pulmonary lesions or granulomas that exhibit caseous necrotic pulmonary lesions with substantial bacterial burden following *M.tb* infection, both intracellularly and extracellularly, closely resembling those seen in human patients [19]. These cavities serve as sites for rapid bacterial multiplication and have direct access to airways, facilitating TB transmission [20] and demonstrate a range of lesion types that mirror various human TB pathologies.

Irwin and colleagues identified three distinct lesion types, namely Type I, Type II, and Type III [19] five weeks after administration of a low-dose aerosol infection. These lesions display differing levels of host control over bacterial replication and inflammatory responses, as shown by morphometric analysis of lesion sizes and bacterial load [19]. Type I lesions closely resemble human TB granulomas, forming solid, encapsulated necrotic caseous lesions, which are characteristic of severe TB pathology. Type II lesions resemble granulocytic pneumonia, characterized by rapid progression and an overwhelming presence of neutrophils, leading to extensive lung consolidation and parenchymal collapse along their periphery. Although Type II lesions share similarities with Type I granulomas, such as neutrophil dominance, they lack the organized fibrotic structure and collagen deposition present in Type I [19]. These Type II lesions in C3HeB/FeJ mice bear resemblance to the polymorphonuclear alveolitis occasionally observed in human TB patients [21]. Finally, Type III lesions are similar to those observed in BALB/c mice following aerosol TB infection but notably lack the caseous necrosis characteristic of other lesion types within the Kramnik model [19].

However, despite its widespread use, the Kramnik model has severe limitations, resulting in notable inconsistencies in the results reported by different research groups, particularly concerning lethality and susceptibility [22,23]. It has a distinct immune profile compared to resistant strains like C57BL/6 or BALB/c mice [24]. The SST1 locus contains various genes implicated in immune functions. One such gene is NRAMP1 (Natural Resistance-Associated Macrophage Protein 1), which is involved in controlling the replication of intracellular pathogens, including mycobacteria [25]. The intracellular pathogen resistance 1 (*Ipr1*) gene within the SST1 locus is involved in the regulation of macrophage apoptosis and necrosis in response to infection – the encoded Ipr1 protein has been shown to play a role in defence mechanisms against intracellular pathogens, and in the cell death pathway of infected macrophages [26,27].

In practice, a significant challenge in working with the Kramnik model is the difficulty in directly sequencing the SST1 locus, which is large and highly redundant [28]. The SST1 locus has been linked to resistance or susceptibility to TB and is significantly associated with their unique response to *M.tb*. Recent studies suggest that sequencing SP110 and particularly SP140 within this locus can assist in determining whether mice are true homozygous Kramniks or have undergone backcrossing. This is especially pertinent for research conducted in regions where TB is endemic, as breeding programs often involve many generations, increasing the risk of backcrossing significantly after the 10th generation [3,29,30].

The immune responses of Kramnik mice also differ from human immune responses, particularly in the representation of macrophage functionality and B-cell responses, which can influence experimental outcomes and complicate the interpretation of results [31]. Notably, while constraints in macrophage pathology are documented [32], the impact on B-cell activity remains insufficiently demonstrated. In addition, the immune system-related deficiencies necessitate a careful approach when extrapolating findings to human scenarios and the immune discrepancies raise important questions about the translational potential of findings derived from the Kramnik model. On the other hand, the interactions between infected mycobacteria and therapeutic molecules show less reliance on the immune environment of the host, especially if the molecules can be delivered intracellularly. The model may therefore be very useful for therapeutic studies. Nevertheless, the Kramnik mouse model remains a critical tool for evaluating novel therapeutic strategies, such as those explored in this study.

### The Pheroid^®^ drug delivery system

Innovative therapeutic delivery methods would be of benefit in addressing the challenges associated with traditional drug administration, particularly in complex diseases such as TB. The Pheroid^®^ drug delivery system represents a significant advancement, enhancing the bioavailability and therapeutic effectiveness of encapsulated agents [33–35]. The flexibility of the Pheroid^®^ system allows it to be tailored to specific applications through modifications in component ratios and preparation methods, making it adaptable to a variety of therapeutic needs. This proprietary formulation utilizes lipid-based carriers to facilitate the efficient transport of antimycobacterial compounds through biological membranes. By creating a stable microencapsulated environment, the Pheroid^®^ system not only improves the solubility of hydrophobic drugs but also enables controlled release and targeted delivery to infected tissues (proprietary information; AF Grobler). These properties make it a promising candidate for integration with standard anti-TB treatment protocols, offering potential improvements in therapeutic outcomes when tested within the Kramnik mouse model in this study.

The Pheroid^®^ system is composed of mostly natural, non-toxic ingredients, including polyunsaturated fatty acids (PUFAs) such as omega-3 and omega-6, known for their minimal side effects [33,35]. This system is economically produced using an environmentally friendly method and can effectively entrap, transfer, and deliver compounds across physiological barriers such as cellular membranes. Structurally, the Pheroid^®^ is a three-phase system comprising a water-phase, an oil-phase (pro-Pheroid^®^), and a gas-phase (nitrous oxide) [33,35,36]. The oil phase, known as pro-Pheroid^®^, contains a mixture of esters of linoleic acid (35.5%), linolenic acid (30.5%), and oleic acid (21.4%). These PUFAs have demonstrated potential in enhancing the delivery of various compounds and exhibit antibacterial activity against *M.tb* [37–39].

### Purpose of the study

#### Evaluating the usability of C3HeB/FeJ mouse model for therapeutic evaluations

**I**n this study, a Pheroid^®^ formulation of the first-line therapeutic drugs RIF, INH, EMB and PZA serves to explore therapeutic responses in the C3HeB/FeJ Kramnik mouse model. In theory, this model may serve as a valuable tool for evaluating therapeutic interventions, but the inconsistencies in its immunological characteristics may pose challenges for translating findings directly to human biology. The variability in immune responses and the presence of unique strain-specific interactions further complicate the extrapolation of data to human TB, underscoring the need for a critical assessment of the model’s strengths and weaknesses.

This study aimed to investigate the usability of the Kramnik mouse model in a therapeutic approach by administering a novel Pheroid^®^ formulation in combination with traditional anti-TB antibiotics. The aim was not to evaluate the formulations as such, but rather to illustrate the model’s possible uses and constraints.

### Materials and methods

#### Materials for treatment formulations

Materials used during the formulation study were the following: vitamin F ethyl ester (Chemimpo, South Africa), Kolliphor^®^ EL (BASF, Germany), DL-α-tocopherol (Chempure Pty. Ltd., South Africa), mycolic acids (MAs) (donated by Dr. Yolandy Lemmer from CSIR, South Africa), Incromega™ E7010-LQ-(LK) (Croda Chemicals SA Pty. Ltd., South Africa) and ascorbyl palmitate (DB Fine Chemicals Pty. Ltd., South Africa). The APIs were EMB, RIF, PZA and INH (DB Fine Chemicals Pty. Ltd., South Africa). Distilled water (Merck Millipore Elix^®^ Essential 3) and medicinal grade nitrogen oxide (N_2_O) (AFROX, South Africa) were also used. While characterising the different formulations, dilutions were made with 32% hydrochloric acid (HCI) (Sigma-Aldrich Pty. Ltd, South Africa), and the Pheroid^®^ vesicles were stained using a 1 mg/ml solution of a Nile Red fluorescent marker (Sigma-Aldrich Pty. Ltd, South Africa).

After adding the different APIs to each part of the formulation, the formulations were first sonicated (Transonic TS 540, USA) and then homogenised (Omni TH_Q_ - Digital Tissue Homogenizer, USA). During the formulation study, the particle size of each formulation was determined with a Malvern Mastersizer Hydro 2000 SM (Malvern Instruments, Worcestershire, United Kingdom), the zeta potential was determined with a Malvern Zetasizer Nano ZS (Malvern Instruments, Worcestershire, United Kingdom), and confocal laser scanning microscopy (CLSM) images were obtained with a Nikon D-eclipse C1 confocal scanning microscope (Nikon, USA).

The Pheroid^®^ system was modified for TB applications by *inter alia* adjusting the ratios of the standard oil-phase ingredients and the addition of eicosapentaenoic acid (EPA) and MAs. EPA, a PUFA with known anti-TB and anti-inflammatory properties, was included in concentrations higher than those used in previous formulations to enhance the system’s therapeutic efficacy [38,40,41]. Additionally, ascorbyl palmitate, an antioxidant known for its ability to prolong the shelf life of oils and reduce oxidative stress, was incorporated to improve the stability and safety of the formulation [42–44].

MAs, unique lipid components of the Mycobacterium cell wall, play crucial roles in the survival, virulence, and growth of *M.tb* [45–47]. In this study, MAs were integrated into the Pheroid^®^ delivery system to potentially act as targeting ligands, directing the therapeutic compounds specifically to infected macrophage cells [48]. By refining these components, the study aimed to leverage the Pheroid^®^ system to enhance the efficacy of anti-TB drugs, reduce toxicity, and improve overall treatment outcomes in the Kramnik mouse model.

#### Materials for the animal study

Two H37Rv mouse passaged *M.tb* strain aliquots were used for infection and were obtained from the Welcome Centre for Infectious Diseases Research in Africa (CIDRI-Africa) and the Institute of Infectious Diseases and Molecular medicine (IDM), University of Cape Town, South Africa. Phosphate buffered saline (PBS), pH 7.2 (Diagnostic Media Products at National Health Laboratory Service (NHLS), South Africa), was used to dilute the *M.tb* for infection. Before infection, the mice were anesthetized using a 2.5% isoflurane (Isofor, Safeline Pharmaceuticals, Johannesburg, South Africa) O_2_/mixture at a rate of 2 L/min. To determine the baseline colony forming units (CFUs) and the bacterial load immediately prior to treatment, harvested lungs were homogenised by means of a QIAGEN TissueLyser II at a frequency of 25/s for 12 min in sterile PBS and plated on Middlebrook 7H11+OADC agar plates (Diagnostic Media Products at NHLS, South Africa) and incubated at 37±2°C in a 5% CO_2_ atmosphere for 3 weeks.

For CFU and bacterial load analysis, the lungs of the mice were sent to CIDRI-Africa and IDM, University of Cape Town, South Africa. Harvested lungs were homogenised using a motorised homogeniser with dispersing tool (Heidolph DIAX 900) in sterile PBS. Serial dilutions (10-fold) were cultivated on Difco^TM^ Middlebrook 7H10 agar plates (BD Biosciences, San Jose, CA), supplemented with 10% Middlebrook OADC, and 0.5% glycerol in triplicate for 3 weeks to evaluate *M.tb* growth. After 3 weeks (21 days) of incubation at 37°C in the dark, the CFUs were counted [49,50].

### Methods

#### Study design and experimental groups

The study employed a structured experimental design to investigate the therapeutic efficacy of the Pheroid^®^ formulation. The anti-TB activity was determined against a standardized dose of H37Rv *M.tb* strain in infected C3HeB/FeJ female mice (8-10 weeks old). The study was approved by the North-West University AnimCare committee of South Africa (Ethics number: NWU-00292-17-A5). The Vivarium of the North-West University, South Africa provided the female C3HeB/FeJ mice (8-10 weeks of age) for this study. This facility is accredited by the Association for Assessment and Accreditation of Laboratory Care International (AAALAC) and is a South African National Accreditation System (SANAS) Good Laboratory Practice (GLP) certified facility. The principles of the 3Rs—Replacement, Reduction, and Refinement were applied during this study.

One hundred and eighty-six mice were divided into four treatment groups. All mice were infected with *M.tb.* The treatment of each of the groups are as follows:

1. Positive control or reference group: received the first-line TB treatment APIs (RIF, INH, PZA, and EMB).
2. Carrier group: received the Pheroid^®^ formulation without APIs.
3. Test formulation group: received the Pheroid^®^ formulation with the first-line TB treatment APIs entrapped.
4. Negative control group: received saline only.

#### Animal study proper

The mice were housed in a Bio-safety level 3 containment unit in enriched individually ventilated cages (IVC) on a rack isolator system. This system controlled the temperature and humidity with 20 air changes per hour while maintaining positive pressure. Room temperature was maintained at 21±2°C and the humidity at 55±10%. A day-night cycle of 12 hours was firmly upheld; food and water were provided *ad libitum*. Bedding (derived from dust free, sterile corncob chips) contributed to a reduction of toxic gasses and a safe environment for the test animals. The animals were subjected to standard health monitoring with body mass measurements and overall health inspection [51]. The mice were observed daily for changes in body weight, behaviour and any other signs of distress or mortality.

#### Infection

A frozen aliquot containing 1 mL of 6.8 x 10^7^ CFU of *M.tb* H37Rv was thawed and diluted 100-fold with PBS. To infect the mice, the animals were anaesthetized by exposing each to a 2.5% isoflurane/O_2_ mixture at a rate of 2 L/min in a 1 L chamber, until properly anesthetized. The animals were closely monitored during the procedure for signs of distress. Each unconscious mouse was removed from the chamber and 50 µL of the 100-fold dilution containing an estimated concentration of 3.4 x 10^4^ CFU of *M.tb* was gradually released into the nostrils with a sterile 20 gauge feeding needle for intranasal infection. The rate of release was adjusted to allow the mouse to inhale the inoculum. The animals were infected in a randomised manner [52]. To verify the actual inoculum titre, serial dilutions of the 100-fold diluted *M.tb* suspension were cultivated in triplicate on 7H11 agar plates supplemented with OADC; CFU were counted after incubation for 3 to 4 weeks at 37°C and 5% atmospheric CO_2_.

To confirm that the infection had indeed been established in the animals, three untreated but *M.tb* infected animals were euthanised by cervical dislocation, two days after infection. Lungs, livers, and spleens were harvested to determine the baseline CFUs. Each organ was weighed and frozen (-22°C) in PBS until analysis. After three weeks, an additional three mice were euthanized to determine the bacterial infection load before treatment was started. Mice were euthanized on the predetermined euthanasia dates, and the lungs were used for CFU analysis.

### Treatment

#### Formulation selection and characterization

Pheroid^®^ formulations with varying oil-to-water *w/w* ratios (1:2, 1:3, 1:5) were prepared and characterized. Formulation characterization was performed using CLSM for visualizing structural morphology and distribution at the cellular level, particle size analyses and zeta potential measurements to assess stability and electrostatic interactions, and particle size distribution analysis to predict pharmacokinetic profiles (Fig 1). The 1:2 formulation, demonstrating superior stability and effective mycobactericidal fatty acid concentrations, was selected for further use in the study.

**Fig 1.**
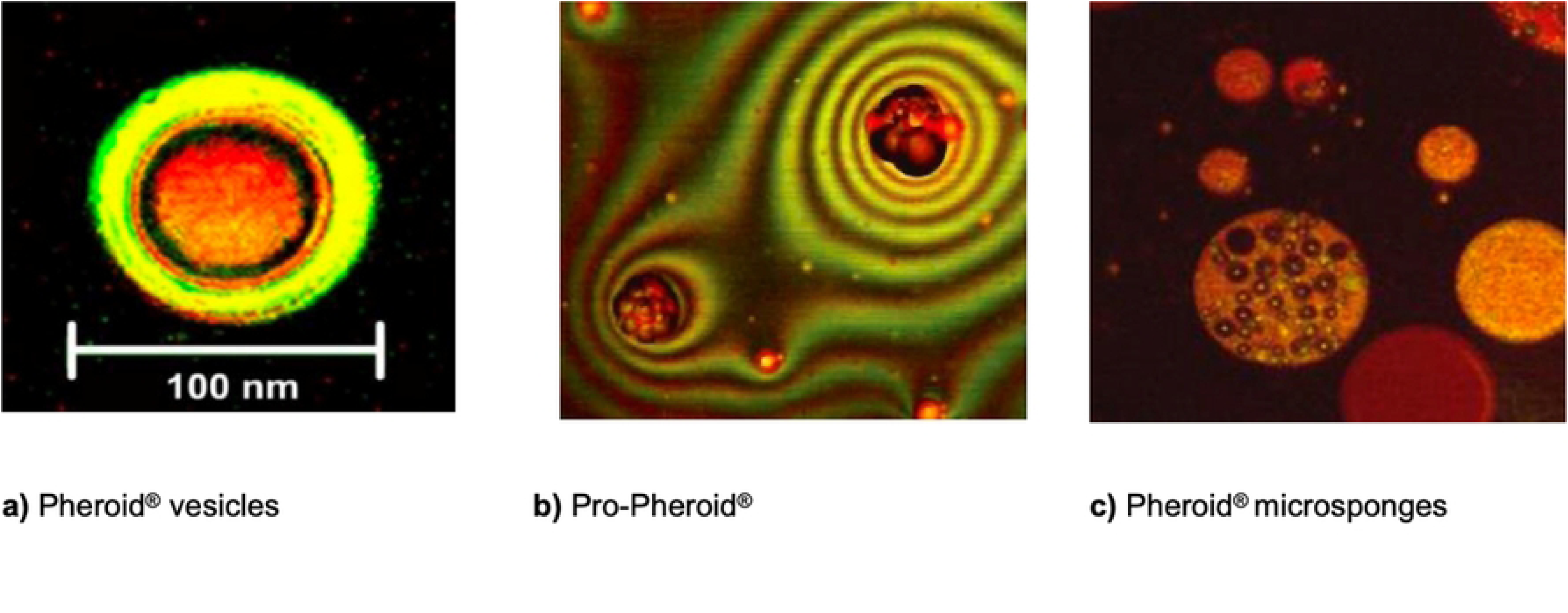
The morphology of different Pheroid^®^ types. (A) Pheroid^®^ vesicles; (B) Pro-Pheroid^®^, and (C) Pheroid^®^ microsponges. (Reprinted from Grobler (2009) [33], with permission from the author).

To evaluate the therapeutic efficacy of the novel Pheroid^®^ formulation, the female C3HeB/FeJ mice were divided into the predefined experimental groups for treatment, which included the standard anti-TB antibiotics and the Pheroid^®^ formulation combined with these antibiotics after confirmation of infection. The study consisted of a treatment phase (3-month duration), and a follow-up phase (3-month duration). During the treatment phase, the various treatments were administered to the mice starting three weeks after infection, where after each test group received the relevant test formulation (150 µL) *via* oral gavage daily, five days a week for 12 weeks. This translated into a dose of 10 mg/kg RIF, 100 mg/kg EMB, 25 mg/kg INH, 150 mg/kg PZA, 0.1 mg/kg ascorbyl palmitate, 1.1 mg/kg MAs and 70 mg/kg EPA [53–55]. During oral gavage, 1 mL sterile pyrogen free syringes (Surgi Plus Medical, Durban, South Africa) and 22 gauge feeding needles (Instech Laboratories Inc, Pennsylvania, USA) were used to administer the treatment formulations.

Animals were humanely terminated at a predefined endpoint of the study in the following manner: six animals from the positive control, carrier and test formulation groups were euthanised at different time intervals, *i.e*. at week 1, 2, 3, 4, 8 and lastly at week 12, and four animals from the negative control group at different time intervals, *i.e*. at week 1, 4 and 8 in accordance with the replace principle of the 3R’s of ethics in animal research.

For the follow-up phase, six animals from the positive control, carrier and test formulation groups were euthanised at different time intervals, *i.e*. at month 1, 2, and 3. The lungs, liver and spleen of each mouse were harvested to provide information regarding disease progression during the treatment phase [20,53]. Inflammation in the lungs indicates the presence of the *M.tb* infection, and a direct correlation can be drawn between the mass of the lungs and the severity of the infection [56]. The lungs were also cultured and the amount of CFUs in the lungs were determined.

### Outcome measurements and assessment criteria

Evaluating the therapeutic efficacy of the Pheroid^®^ formulation in the Kramnik mouse model included organ mass indices (organ weight/body weight x 100) and CFU counts to assess bacterial load.

### Statistical analysis

To evaluate the therapeutic efficacy of the Pheroid^®^ formulation, the viable bacterial counts of the lung samples were calculated. Descriptive statistics provided an overview of data characteristics, including mean, median, and standard deviation. To quantify any possible effect, the data was expressed as the mean Log 10 CFU ± the standard deviation. The mean organ weight and CFU values were evaluated by one-way analysis of variance (ANOVA), followed by Tukey multiple comparison or Kruskal-Wallis tests. Differences in means were considered significant if p ≤ 0.05 (*i.e.* at the 5% level of significance), using STATISTICA statistical software (StatSoft, Inc., 2018) and Prism GraphPad (2025). ANOVA and post hoc tests identified significant differences among treatment groups while regression analysis was used to evaluate relationships between dosing regimens and observed outcomes. Because no data could be recorded for the carrier control’s month 3, months 1 and 2 were compared to each other by means of a T-test, as reported. A substantial number of mice from the negative control group became *moribund* and had to be euthanized as per study protocol. This resulted in a loss of statistical significance for this group.

## Results

### Therapeutic efficacy of the Pheroid^®^ Formulation

The methodological framework employed in this study was designed to assess both the therapeutic efficacy of the novel Pheroid^®^ formulation as well as the suitability of the well-characterized C3HeB/FeJ Kramnik mouse model of *M.tb* infection for therapeutic interventions. A systematic approach was utilized, beginning with the intranasal inoculation of female mice with a standardized dose of H37Rv strain bacteria. Subsequent randomization into four treatment groups enabled comparative analysis between the Pheroid^®^ formulation combined with first-line anti-TB therapies and control groups. Formulation characterization was performed using *inter alia* CLSM and zeta potential measurements, which confirmed the formulation’s stability and mycobactericidal properties.

The mice were observed daily for changes in body weight, behaviour and any other signs of distress or mortality. Euthanasia at predetermined intervals allowed for comprehensive assessment of organ health through mass index calculations and CFU analysis, thereby elucidating the relationship between treatment and bacterial load reduction. Because the negative control group received no treatment and because of the heterogeneity of this model, the mice included in the negative control group became *moribund* early on in the study and had to be euthanized as per study protocol. No resulting statistical significance could be drawn for the negative control group; however, the data was recorded and given for each organ.

### Formulation characterization

The 1:2 Pheroid^®^ formulation displayed the highest mycobactericidal fatty acid inclusion and the smallest mean particle size, making it the most promising candidate for therapeutic application.

The 1:2 (oil: H_2_O) formulation formed Pheroid^®^ carriers that complied with specifications (depicted in Table 1 and Fig 2); these carriers had a median particle size of 0.196 μm, and 80% of particles distributed between 0.077 μm and 4.371 μm. The zeta potential was measured at -15.3±4.42 mV, indicating stability. Despite incomplete dissolution of APIs, the formulation stayed in suspension for optimal oral delivery.

**Fig 2.**
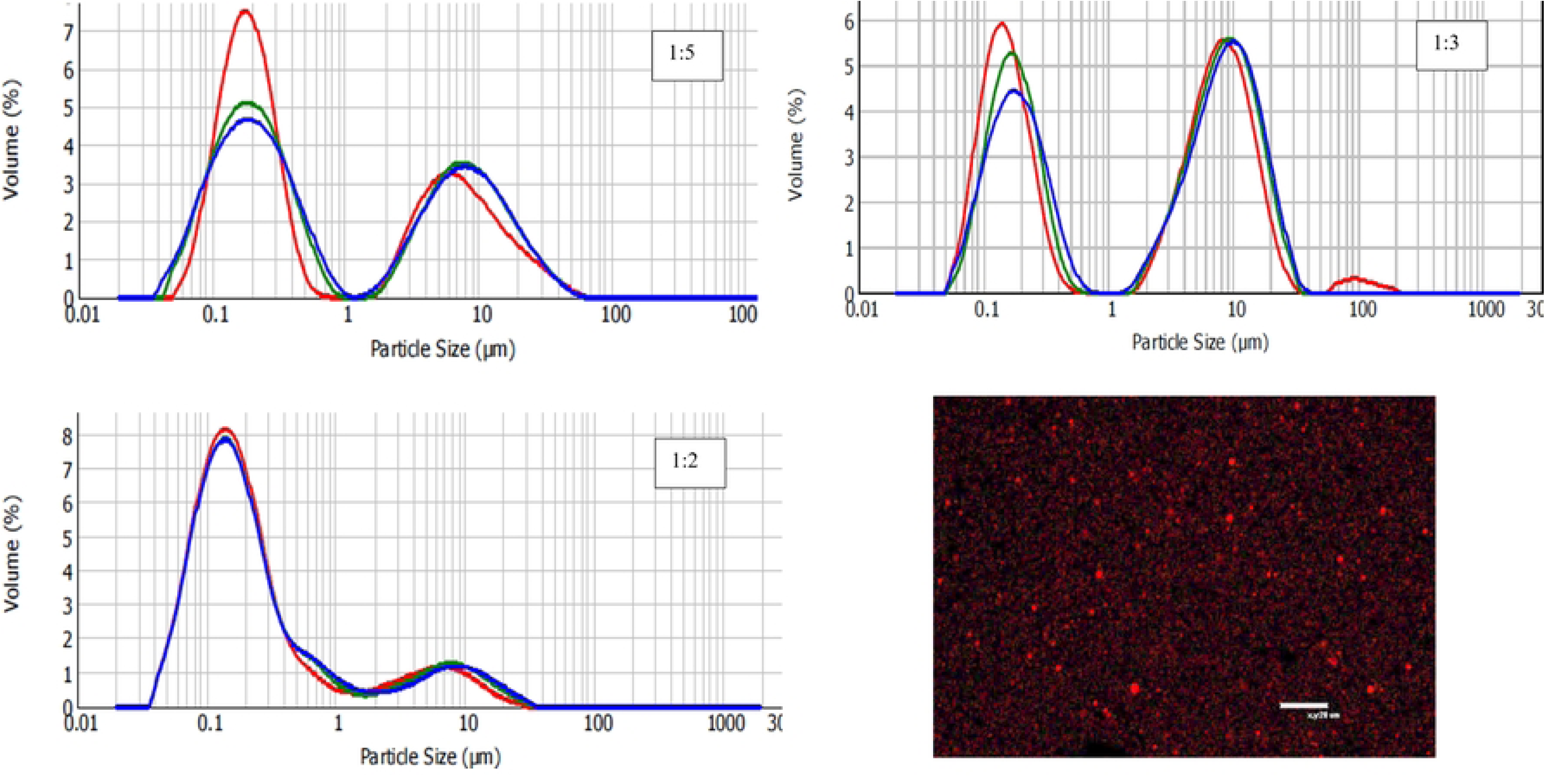
Specifications of Pheroid^®^ carriers. A confocal laser scanning microscopy micrograph depicting the morphology of the Nile red stained Pheroid^®^ carriers and the particle size distribution spectra of the different oil: H_2_O ratios.

**Table 1.** Particle size distribution and Zeta potential of Pheroid^®^ carriers.

| Formulation | Zeta potential (mV) | Particle size ($\mu\text{m}$)<sup>a</sup> |
| --- | --- | --- |
| 1:2 control | $-18.5 \pm 3.76$ | 0.177 |
| 1:2 | $-15.3 \pm 4.42$ | 0.196 |
| 1:3 control | $-18.8 \pm 4.04$ | 0.178 |
| 1:3 | $-20.2 \pm 4.14$ | 4.376 |
| 1:5 control | $-27.0 \pm 6.29$ | 0.433 |
| 1:5 | $-17.0 \pm 3.57$ | 0.352 |

<sup>a</sup>Mean provided for three measurements per sample.

### Survival rates

In evaluating the therapeutic efficacy of the Pheroid^®^ carrier formulation, significant differences in survival rates among the various treatment groups were observed (Fig 3A). The control group, which received no treatment, exhibited a 100% mortality rate within eight weeks post-infection, with 50% of the mice having to be humanely terminated within the first two weeks after infection, and only one mouse surviving up to week eight, highlighting the virulent nature of H37Rv in the absence of intervention. Conversely, the positive control group receiving standard anti-TB therapies achieved complete survival, demonstrating the effectiveness of pharmacological intervention for this bacterial strain.

**Fig 3.**
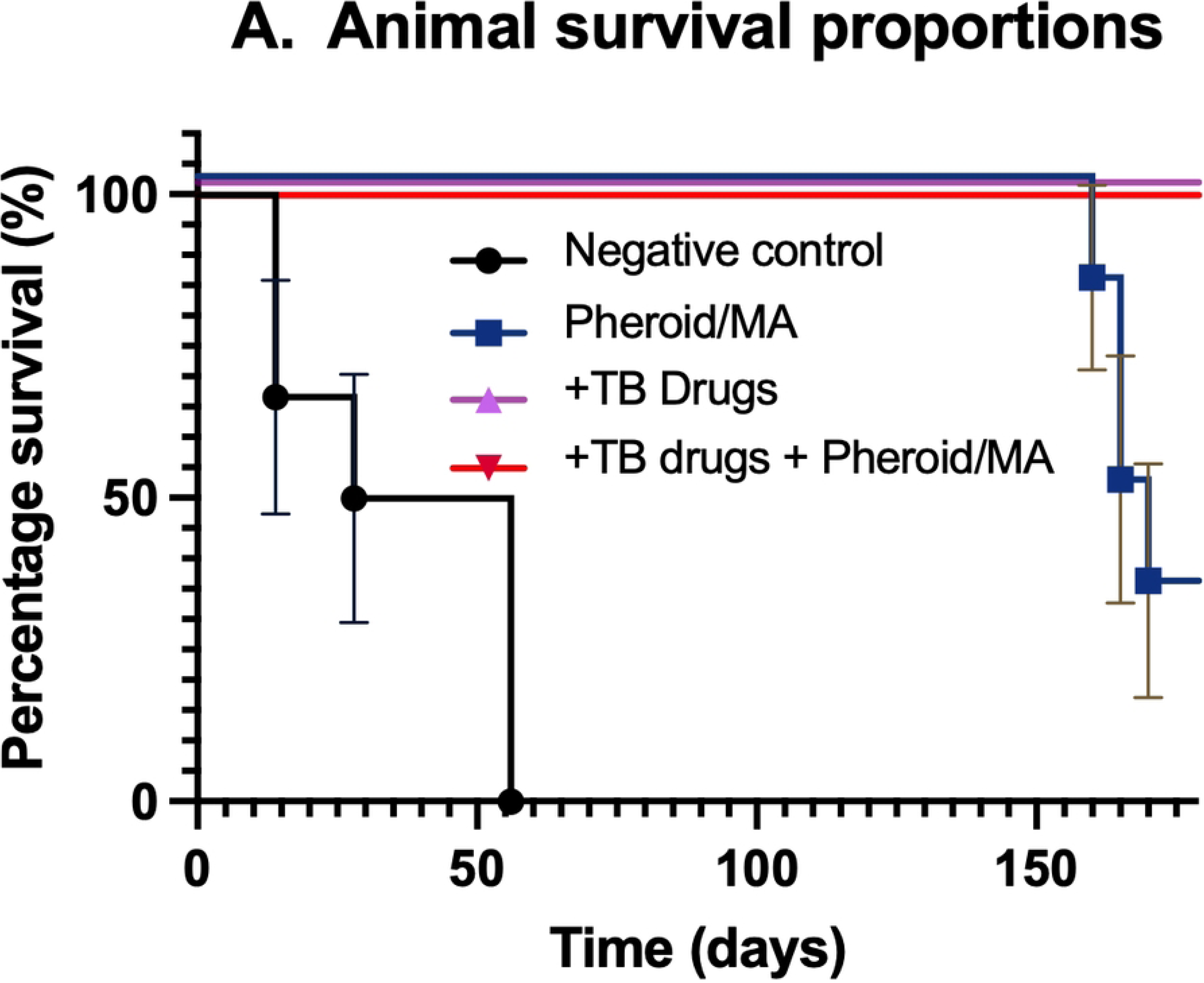
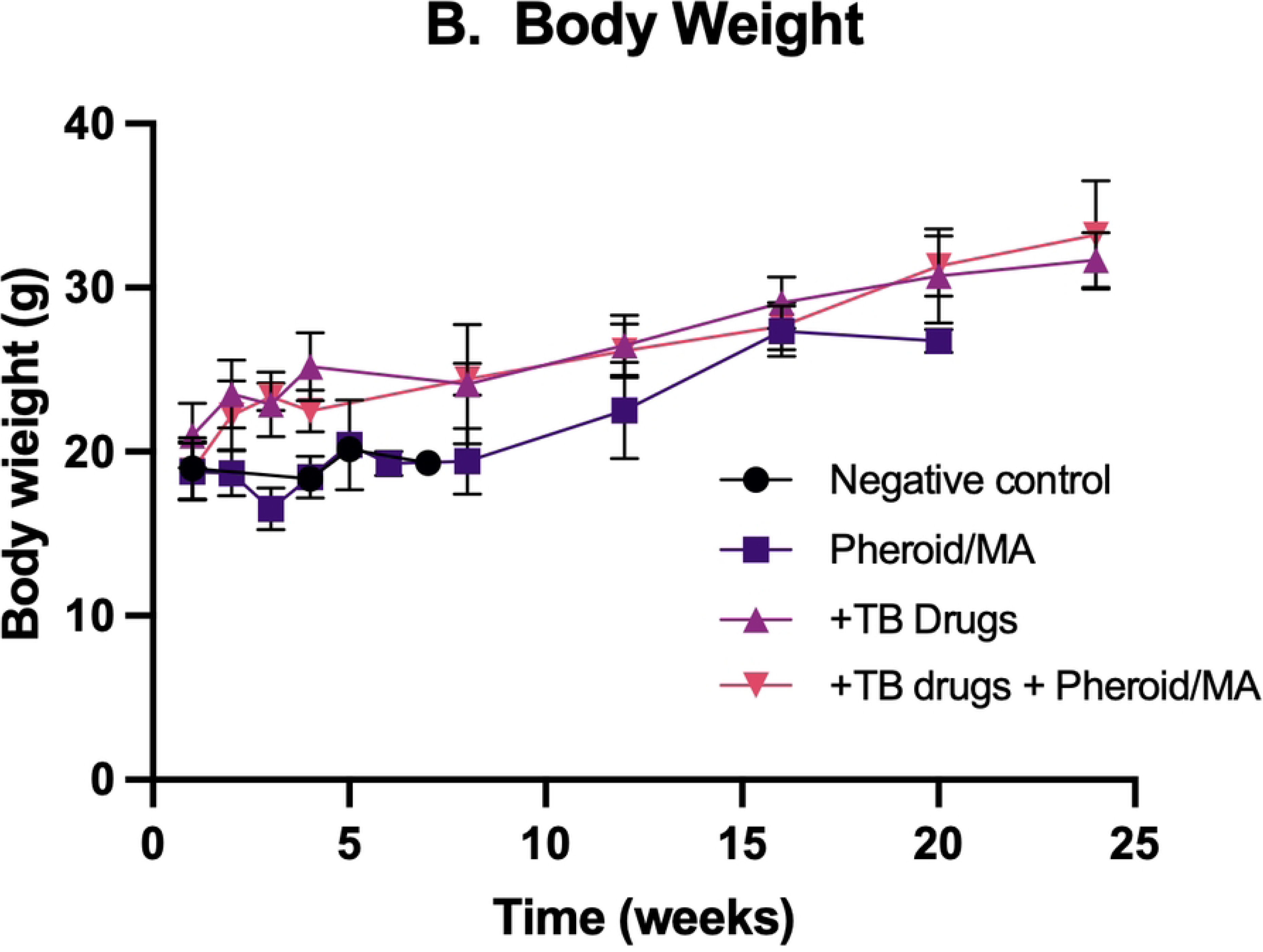
Survival rates and body weight of test animals. (A) Survival of animals after infection and therapeutic intervention. (B) Body weight of the animals after infection and during therapy by oral gavage over the course of the study. The control group received a similar amount of water by oral gavage.

No animal in the carrier group died, even though no antibiotics were entrapped in the carrier. However, as can be observed in Fig 3B, the animals in that group initially lost weight, whereas the groups receiving the conventional antibiotic regimen or the group receiving the antibiotic regimen entrapped in the carrier started gaining weight immediately after initiation of therapy and continued gaining weight over the course of the study. The animals in the carrier group also started gaining weight from week three of the intervention. The carrier group still showed a lower mean body weight of nearly 15% than the test formulation and positive control groups. The negative control showed no significant weight gain.

### Lung, liver, and spleen indices

The lungs, liver and spleen of each mouse in the prospective groups *i.e*. positive control containing the conventional TB antibiotic regimen, the carrier group, the test formulation containing the conventional TB antibiotic regimen entrapped in the carrier and negative control, were harvested during the study as from week three of the study. The mass index of each organ, *i.e.* (organ weight/total animal weight) x 100, and the *M.tb* CFUs present in each harvested lung were determined. Organ indices were measured to assess the impact of both the TB infection and the efficacy of therapeutic interventions. The indices determined in this study were the lung, liver, and the spleen index.

In TB infection, infected mice may display an elevated lung index due to disease-induced hypertrophy and granuloma formation. An increased lung index may therefore indicate a higher disease burden, with more pronounced lung pathology, including granulomas, inflammation, and tissue damage. Successful therapeutic interventions will in such cases show a reduction in the lung index compared to untreated control, since that would also correspond with a decrease in lung inflammation and pathology [57,58].

A comparison of the mean organ indices over time is presented in Table 2 and visually in Fig 4. In this study, a comparison of the lung indices throughout the treatment phase, *i.e*. weeks 3, 4, 8 and 12, of the positive control *vs.* carrier *vs.* test formulation, showed that the lung index of the positive control and test formulation differed significantly (p ≤ 0.05) from the carrier group at each time point, providing a clear indication that both the positive control and the test formulation suppressed the *M.tb* infection to the same extent and much better than the carrier only. The positive control and test formulation did not differ in a statistically significant manner from one another.

**Fig 4.**
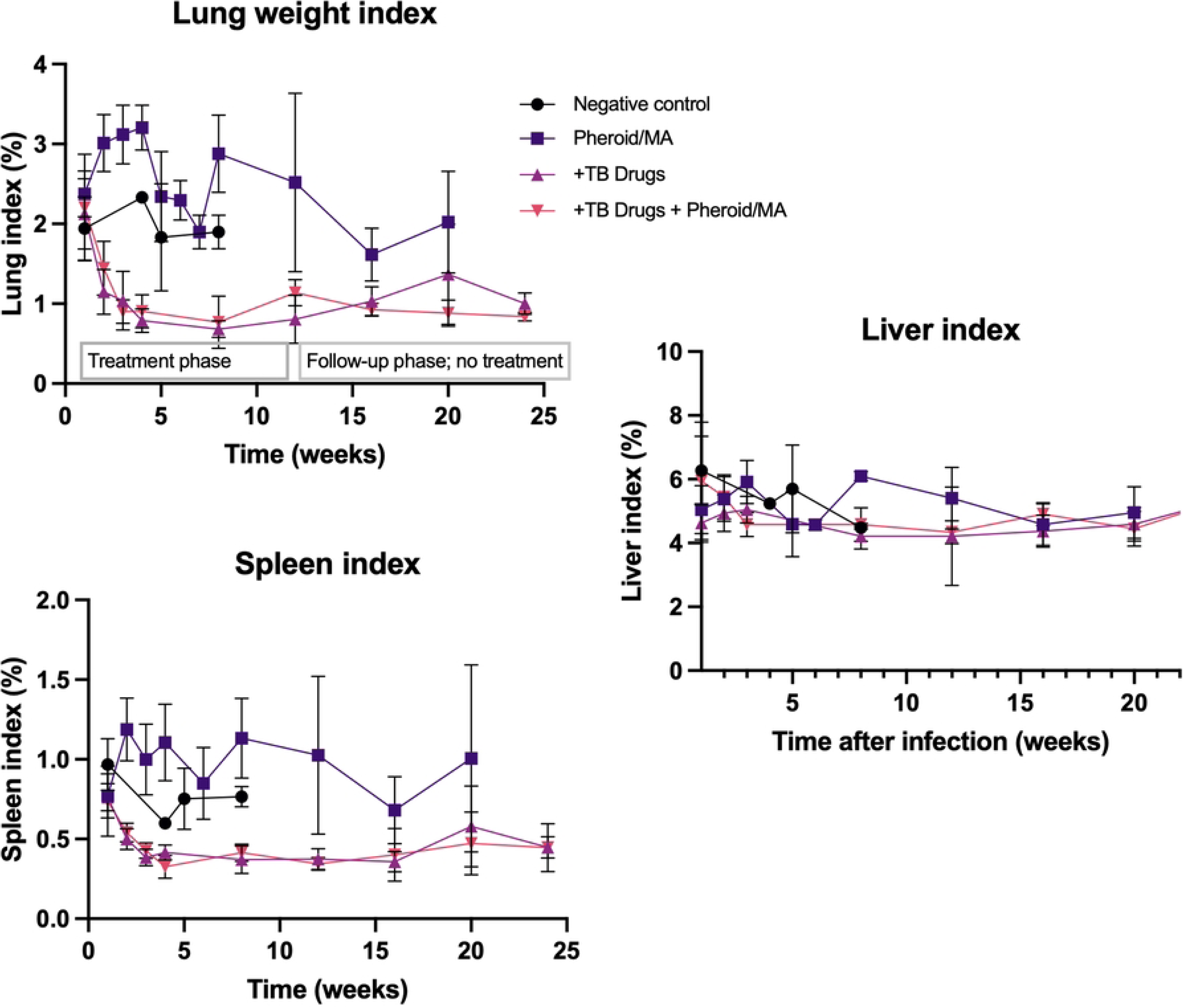
Organ indices in Kramnik mice during treatment with different therapeutic interventions.

**Table 2.**
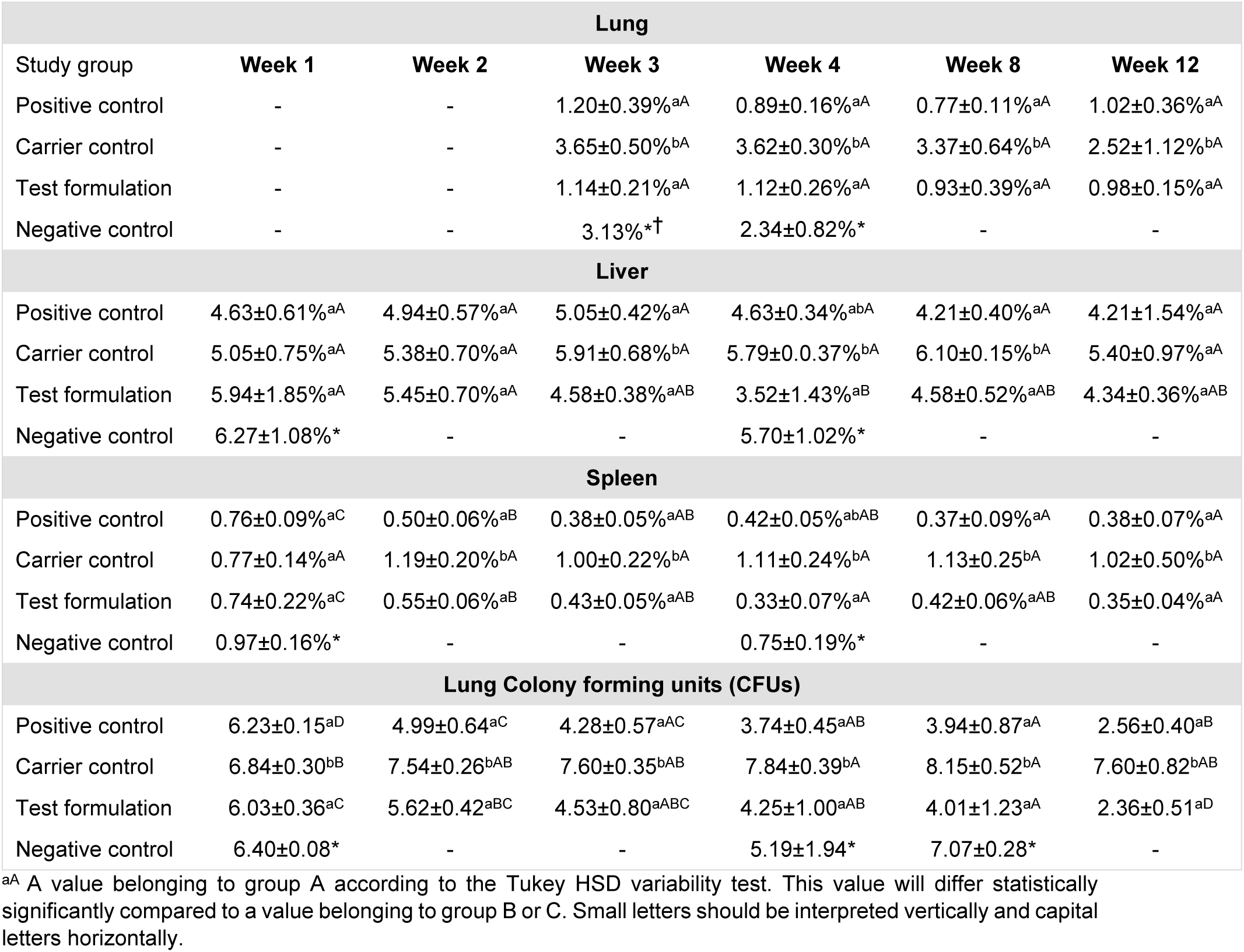
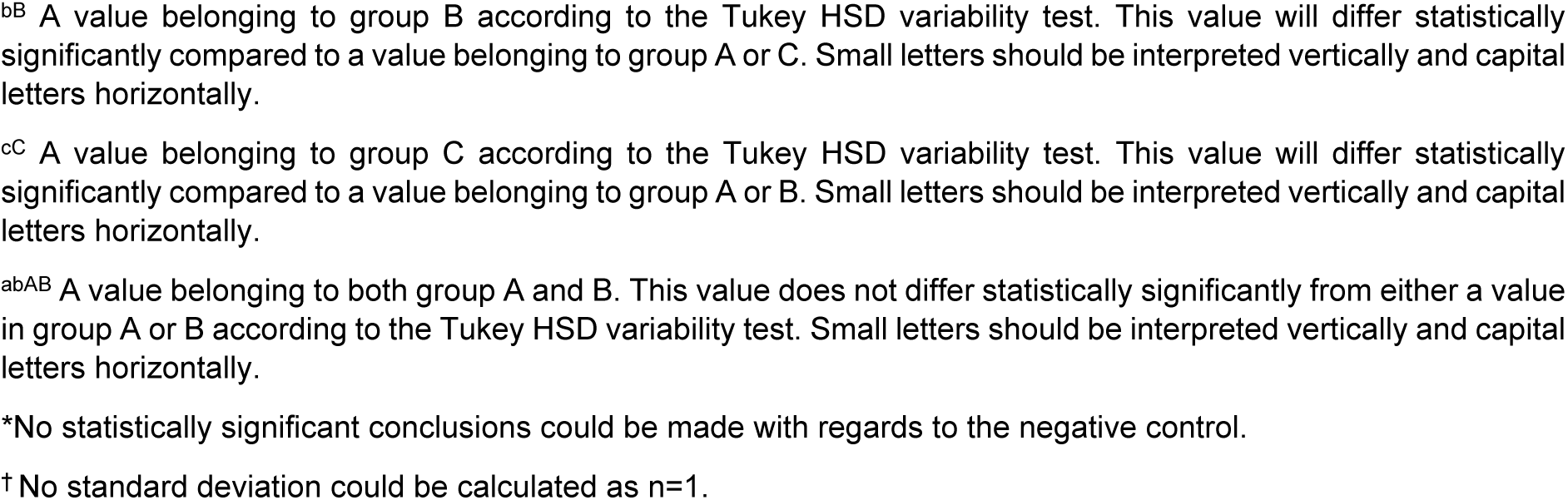
Measured indices during the treatment phase.

| Lung |  |  |  |  |  |  |
| --- | --- | --- | --- | --- | --- | --- |
| Study group | Week 1 | Week 2 | Week 3 | Week 4 | Week 8 | Week 12 |
| Positive control | - | - | 1.20±0.39%<sup>aA</sup> | 0.89±0.16%<sup>aA</sup> | 0.77±0.11%<sup>aA</sup> | 1.02±0.36%<sup>aA</sup> |
| Carrier control | - | - | 3.65±0.50%<sup>bA</sup> | 3.62±0.30%<sup>bA</sup> | 3.37±0.64%<sup>bA</sup> | 2.52±1.12%<sup>bA</sup> |
| Test formulation | - | - | 1.14±0.21%<sup>aA</sup> | 1.12±0.26%<sup>aA</sup> | 0.93±0.39%<sup>aA</sup> | 0.98±0.15%<sup>aA</sup> |
| Negative control | - | - | 3.13%*<sup>†</sup> | 2.34±0.82%* | - | - |
| Liver |  |  |  |  |  |  |
| Positive control | 4.63±0.61%<sup>aA</sup> | 4.94±0.57%<sup>aA</sup> | 5.05±0.42%<sup>aA</sup> | 4.63±0.34%<sup>abA</sup> | 4.21±0.40%<sup>aA</sup> | 4.21±1.54%<sup>aA</sup> |
| Carrier control | 5.05±0.75%<sup>aA</sup> | 5.38±0.70%<sup>aA</sup> | 5.91±0.68%<sup>bA</sup> | 5.79±0.37%<sup>bA</sup> | 6.10±0.15%<sup>bA</sup> | 5.40±0.97%<sup>aA</sup> |
| Test formulation | 5.94±1.85%<sup>aA</sup> | 5.45±0.70%<sup>aA</sup> | 4.58±0.38%<sup>aAB</sup> | 3.52±1.43%<sup>aB</sup> | 4.58±0.52%<sup>aAB</sup> | 4.34±0.36%<sup>aAB</sup> |
| Negative control | 6.27±1.08%* | - | - | 5.70±1.02%* | - | - |
| Spleen |  |  |  |  |  |  |
| Positive control | 0.76±0.09%<sup>aC</sup> | 0.50±0.06%<sup>aB</sup> | 0.38±0.05%<sup>aAB</sup> | 0.42±0.05%<sup>abAB</sup> | 0.37±0.09%<sup>aA</sup> | 0.38±0.07%<sup>aA</sup> |
| Carrier control | 0.77±0.14%<sup>aA</sup> | 1.19±0.20%<sup>bA</sup> | 1.00±0.22%<sup>bA</sup> | 1.11±0.24%<sup>bA</sup> | 1.13±0.25%<sup>bA</sup> | 1.02±0.50%<sup>bA</sup> |
| Test formulation | 0.74±0.22%<sup>aC</sup> | 0.55±0.06%<sup>aB</sup> | 0.43±0.05%<sup>aAB</sup> | 0.33±0.07%<sup>aA</sup> | 0.42±0.06%<sup>aAB</sup> | 0.35±0.04%<sup>aA</sup> |
| Negative control | 0.97±0.16%* | - | - | 0.75±0.19%* | - | - |
| Lung Colony forming units (CFUs) |  |  |  |  |  |  |
| Positive control | 6.23±0.15<sup>aD</sup> | 4.99±0.64<sup>aC</sup> | 4.28±0.57<sup>aAC</sup> | 3.74±0.45<sup>aAB</sup> | 3.94±0.87<sup>aA</sup> | 2.56±0.40<sup>aB</sup> |
| Carrier control | 6.84±0.30<sup>bB</sup> | 7.54±0.26<sup>bAB</sup> | 7.60±0.35<sup>bAB</sup> | 7.84±0.39<sup>bA</sup> | 8.15±0.52<sup>bA</sup> | 7.60±0.82<sup>bAB</sup> |
| Test formulation | 6.03±0.36<sup>aC</sup> | 5.62±0.42<sup>aBC</sup> | 4.53±0.80<sup>aABC</sup> | 4.25±1.00<sup>aAB</sup> | 4.01±1.23<sup>aA</sup> | 2.36±0.51<sup>aD</sup> |
| Negative control | 6.40±0.08* | - | - | 5.19±1.94* | 7.07±0.28* | - |

<sup>aA</sup> A value belonging to group A according to the Tukey HSD variability test. This value will differ statistically significantly compared to a value belonging to group B or C. Small letters should be interpreted vertically and capital letters horizontally.
<sup>bB</sup> A value belonging to group B according to the Tukey HSD variability test. This value will differ statistically significantly compared to a value belonging to group A or C. Small letters should be interpreted vertically and capital letters horizontally.
<sup>cC</sup> A value belonging to group C according to the Tukey HSD variability test. This value will differ statistically significantly compared to a value belonging to group A or B. Small letters should be interpreted vertically and capital letters horizontally.
<sup>abAB</sup> A value belonging to both group A and B. This value does not differ statistically significantly from either a value in group A or B according to the Tukey HSD variability test. Small letters should be interpreted vertically and capital letters horizontally.
\*No statistically significant conclusions could be made with regards to the negative control.
<sup>†</sup> No standard deviation could be calculated as n=1.

Comparing the lung index for each group, *i.e*. positive control, carrier and test formulation, between the relevant time points, *i.e.* week 3 *vs.* 4 *vs.* 8 *vs.* 12, indicates that the *M.tb* suppression activity of each treatment group (or the lack thereof), has mostly been achieved during the first three weeks of treatment, with little change thereafter. Data could only be collected for the negative control group for week 3 (n=1) and week 5 (n=1).

TB infection may result in hepatomegaly because of hepatic infiltration by immune cells, liver damage / side effects caused by the antibiotics used in treatment or may even be due to hepatic granuloma formation [59–61]. Conversely, a decrease in the liver index following treatment may be caused by reduced systemic impact or improved liver condition through effective therapy. A comparison of the liver indices throughout the treatment phase, *i.e*. weeks 1, 2, 3, 4, 8 and 12, of the positive control *vs.* carrier *vs.* test formulation, showed that the positive control, carrier group and test formulation group did not show a statistically significant difference (p > 0.05) from one another during weeks 1, 2, and 12 (Table 2). However, during weeks 3, 4 and 8, the test formulation had statistically significantly (p ≤ 0.05) lower liver indices compared to the carrier group. The positive control behaved similarly to the test formulation during weeks 3 and 8.

Comparing the liver indices for each group, *i.e*. positive control, carrier control and test formulation, between the relevant time points, *i.e.* week 1 *vs.* 2 *vs.* 3 *vs.* 4 *vs.* 8 *vs.* 12, the positive control and carrier showed no statistically significant difference between the different time points. However, the test formulation showed a statistically significant decrease in the liver index for weeks 1 and 2 to week 4, whilst no difference could be observed for weeks 3, 8 or 12 (Table 2). Data could only be collected for the negative control group for weeks 1 and 5. A comparison of the liver mean index for each group over time is visually represented in Fig 4.

Spleen hypertrophy may be due to an immune response to TB infection, with increased lymphoid proliferation or sequestration of mycobacteria [58]. Effective treatment resulting in a reduction in the TB burden may lead to a decreased spleen index. Comparing the spleen mean indices throughout the treatment phase, *i.e*. weeks 1, 2, 3, 4, 8 and 12, of the positive control *vs.* carrier control *vs.* test formulation, at each individual time point, the positive control, carrier control and test formulation did not show a statistically significant difference (p > 0.05) from one another during week 1 (Table 2). However, from week 2 and onwards the positive control and the test formulation had statistically significantly (p ≤ 0.05) lower spleen indices compared to the carrier group. The only exception was during week 4, where the positive control did not differ significantly from either the carrier control or the test formulation. When comparing the spleen indices for each group between the relevant time points, the positive control showed a statistically significant decrease for weeks 1, 2, 3, 4, 8 and 12. Week 2 also showed a statistically significant difference from weeks 1, 8 and 12. The carrier control showed no statistically significant difference when comparing any of the weeks to one another. The test formulation showed a statistically significant decrease in the spleen index when week 1 is compared with any other week. Week 2 also showed a statistically significant difference from weeks 4 and 12. Data could only be collected for the negative control group for weeks 1 and 5. A comparison of the spleen mean index for each group over time is visually represented in Fig 4.

In summary, the test and positive control formulations showed similar results in terms of lung, liver, and spleen indices, indicating effective suppression of disease progression. The carrier control formulation was less effective than the test and positive control formulations, except in the liver index during specific weeks.

### Colony forming unit (CFU) analysis

CFU analysis over time served as a critical metric for evaluating the efficacy of the therapeutic interventions. Following initial intranasal inoculation with 34,000 CFUs, significant declines in CFU counts were observed in groups treated with the Pheroid^®^ formulation combined with standard anti-TB antibiotics (Table 2). By week 1, both the positive control and test formulation groups exhibited substantial reductions in bacterial loads, indicating the initial effectiveness of the formulation against *M.tb*, while the negative control group faced complete mortality within four weeks. Analysis of lung CFUs at weeks 4, 8, and 12 showed the carrier control group had significantly higher CFU counts compared to both the positive control and test formulation groups. Notably, the test formulation and positive control groups did not differ significantly from each other at any time point, suggesting comparable efficacy.

A comparison of the CFUs of the lungs harvested for each group, *i.e.* positive control, carrier and test formulation, between the relevant time points, *i.e.* week 1 *vs.* 2 *vs.* 3 *vs.* 4 *vs.* 8 *vs.* 12, showed that the positive control group was significantly decreased from week 1 to week 12. Week 8 had statistically significantly lower CFUs compared to weeks 1 and 2 and had statistically significantly higher CFUs compared to week 12. Week 12 had a statistically significantly lower CFU value compared to all other weeks, except week 4 (p≤0.05). The carrier group showed a statistically significant increase in CFUs when week 1 was compared with weeks 4 and 8, but showed no difference between weeks 2, 3 and 12. The Pheroid^®^ test formulation group showed a statistically significant decrease in CFUs when comparing week 1 with weeks 4, 8 and 12. Week 12 had a statistically lower CFU count compared to all the other weeks. For both the positive control and the test formulation groups there was a statistically significant decline in the number of CFUs when comparing week 1 with week 12. The carrier control however, had no statistically significant decline in the number of CFUs comparing week 1 with week 12. Data could only be collected for the negative control group for week 1 (6.40±0.08), and 4 (5.19±1.94), with only one mouse remaining at week 8. Because of the small sample size and irregular euthanasia intervals, no statistical analysis could be conducted on the negative control. The statistical analyses can be found in Table 2. Fig 5 is a visual representation of the CFU counts for the lungs for each group over the treatment phase.

**Fig 5.**
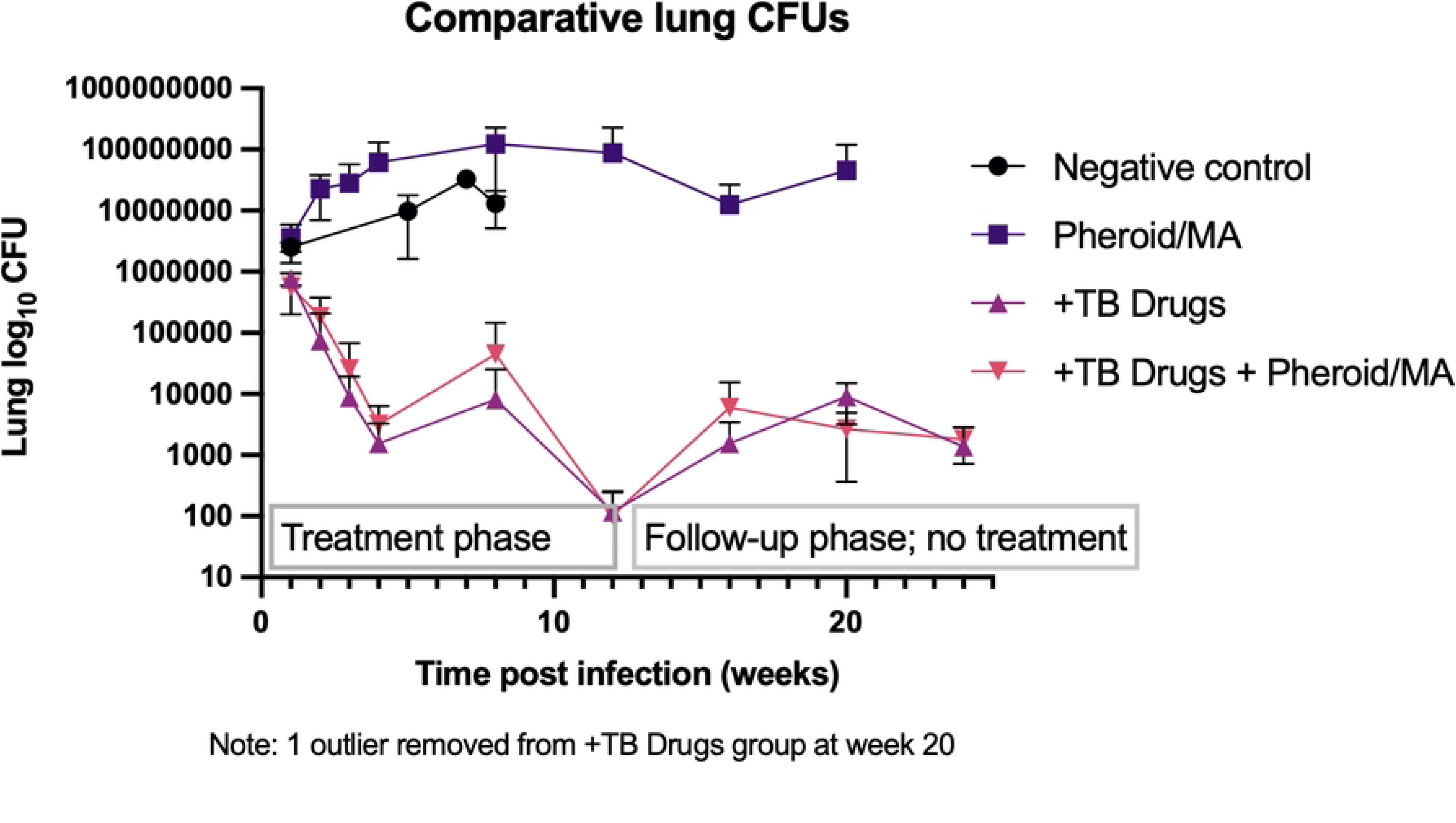
The average lung CFU / gram sample (log10) at different time points during the treatment and follow-up phases. A summary of the CFUs in the study shows the following: Initial inoculum: Mice were infected with an average of 34,000 CFUs intranasally; Week 1: Significant reduction in CFU counts for the test formulation and positive control groups compared to the carrier control and negative control groups; Week 2: Further reduction in CFU counts in the test formulation and positive control groups, while the negative control group had a 100% mortality rate; Weeks 3-12: The test formulation group and positive control group maintained a low CFU count, demonstrating long-term suppression of bacterial load. The CFUs in the Pheroid^®^/MA carrier group were slightly higher than that of the negative control group, yet despite the high titer the animals did not die during the treatment phase; Weeks 12-24. The test formulation group and positive control group maintained a low CFU count, demonstrating long-term suppression of bacterial load even though treatment was discontinued. Mice from the Pheroid^®^/MA carrier group became *moribund* during the follow-up phase, when treatment was discontinued.

Large inter-individual variations in lung CFU counts of specifically the Kramnik mouse model following low-dose aerosol infection have been described, which ultimately causes a reduction in the power of discriminating differences between treatment groups [20,62,63]. For that reason, the very high dose of 34,000 CFUs were used in this study. Four animals in the negative control group were scheduled to be euthanised at different time intervals, *i.e*. at weeks 1, 8 and 12. However, after week 1, mice had to be euthanised earlier than the scheduled dates as they became *moribund*. The fact that the negative control became *moribund* whilst most animals from the other groups did not, is an indication that (a) the *M.tb* infection was successful and that at the high infection dose used it was sufficient to cause the death of untreated animals and; (b) that the disease in this group was much more profound than the other three treatment groups. Additionally, the average CFUs of the three untreated mice (infected with *M.tb*) that were euthanised two days after infection (baseline CFUs), *i.e.* 3.63±0.74 CFU (log10), showed that the infection was successful. Furthermore, the average of the three mice that were euthanised before treatment started (bacterial infection load) was 6.31±0.81 CFU (log 10), indicating that the infection was well established before treatment was initiated. This increase in *M.tb* bacterial load is common in the TB susceptible Kramnik mouse model [64]. It is, however, noteworthy that when utilising the Kramnik mouse model, a higher mortality rate was to be expected, as shown in a study by Lanoix et al. [55], where the death rate of Kramnik mice compared to BALB/c mice was 29.6% *vs.* 3.6%, respectively with a p < 0.01. Driver and co-workers also noted the high mortality of the Kramnik mouse model: in their study 29% of the control group succumbed to the disease three weeks post aerosol infection [20]. No significant differences were found between the groups in the liver index, used as a measure of toxicity and general health.

Comparing the CFUs throughout the treatment phase, *i.e*. weeks 1, 2, 3, 4, 8 and 12, of the positive control *vs.* Pheroid^®^/MA carrier *vs.* test formulation, the positive control and test formulation showed statistically significant differences from the Pheroid^®^/MA carrier at each time point. The positive control and test formulation did however not differ significantly from one another during either the treatment or follow-up phase. Comparing the CFUs of the lungs harvested for each group during the treatment phase, both the positive control group and the test formulation statistically reduced the CFUs in the lungs, but not the Pheroid^®^ carrier. Discontinuance of treatment led to pronounced disease for the carrier group.

The Pheroid^®^ formulation in combination with the APIs (test formulation) did not reduce the *M.tb* infection to a greater extent than the normal treatment regime (positive control). The Pheroid^®^ formulation on its own (carrier control) reduced the *M.tb* infection to a lesser extent when compared to both the test formulation and the positive control. For the test and the positive control formulations, the biggest decline in the number of CFUs in the lungs were observed between week 1 *vs.* 2 (positive control) and week 1 *vs.* 4 (test formulation) of the treatment phase. Thereafter, the number of CFUs in the lungs for both these two groups remained roughly the same until week 12 of the follow-up phase. In neither of these two groups were the mice completely cleared of the *M.tb* infection.

## Discussion

The lung index as an indication of inflammation in the lungs [65] is a useful metric to track TB disease progression and *M.tb* bacterial load in the lungs. Comparing the lung indices harvested throughout the **treatment** phase, *i.e*. weeks 3, 4, 8 and 12, of the positive control *vs.* carrier *vs.* test formulation, the mean lung index of the carrier formulation differed statistically significantly from the mean lung index of the positive control and the test formulation, whereas the positive control and the test formulation did not differ statistically significantly from one another - both the positive control and the test formulation suppressed the *M.tb* infection much better than the carrier formulation. However, the test formulation did not suppress the *M.tb* infection to a greater extent than the positive control. According to the lung index, the *M.tb* disease progression did not change for either of the positive control or test groups after the first three weeks of treatment. For the negative control group data was collected for weeks 3, 4 and 8 only. No statistical analysis could be performed on this group due to early mortality.

Comparing the lung indices harvested throughout the **follow-up phase,** *i.e*. weeks 4, 8 and 12, of the positive control *vs.* carrier *vs.* test formulation, the same trend was observed where the mean lung index of the carrier formulation differed statistically significantly from the mean lung index of the positive control and the test formulation. The positive control and the test formulation did not differ statistically significantly from one another. Comparing the lung index for each group during the follow-up phase, *i.e*. positive control, carrier and test formulation, between the relevant time points, *i.e.* week 4 *vs.* 8 *vs.* 12, no statistical difference was observed between week 4 *vs.* 12 (positive and test formulations) and week 4 *vs.* 8 (carrier control), again indicating there was no change in disease progression for either of the groups during the follow-up phase with regards to the lung index. In this context, the Pheroid^®^ formulation in combination with the APIs (test formulation) did not reduce the *M.tb* infection to a lesser or a greater extent than the normal treatment regime (positive control). Unexpectedly, the Pheroid^®^ formulation on its own (carrier control) reduced the *M.tb* infection to a significant degree compared to the negative control.

In the Lanoix et al. study of 2015, the Kramnik mice model showed remarkable heterogeneity in the timing and extent of disease progression [55], similar to the results of this study. This may explain why the efficacy of both the test formulation and positive control (which both contained APIs) seemed lower than in previous Pheroid^®^ studies done in other models [51,66]. In a study conducted in C57BL/6 inbred mice, Matthee [66] found that by using the Pheroid^®^ delivery system, the RIF plasma concentration could be increased by 305%, subsequently raising the efficacy of the treatment. Nieuwoudt [51] found that the bioavailability of each active ingredient (INH, RIF, EMB and PZA) was enhanced when incorporated into the Pheroid^®^ drug delivery system in comparison with Rifafour^®^ e-275, in female BALB/c mice. In an efficacy study with inbred, specified pathogen-free female BALB/c mice anti-TB drugs (INH, RIF, EMB and PZA) incorporated into the Pheroid^®^ was shown to be more effective with a 10% increase at week 4 and a 16.7% increase at week 12 post-infection [51].

The relationship between plasma drug exposures and antibacterial effect in animal models may be influenced by the differences that can occur in lesion microenvironments, drug penetration into these lesions, the bacterial burden and the ratio of intracellular to extracellular bacilli, which can consequently affect the representation of a certain drug or formulations efficacy, with critical implications for the model in question and its use in forecasting efficacy in humans [55,67]. In a direct comparison of the Kramnik mice model and BALB/c mice using the tuberculosis INH, RIF, PZA, metronidazole and linezolid drugs, the Kramnik mouse model was found to be superior and significantly more refractory to test anti-TB drugs. Although Kramnik mouse models have a less positive treatment outcome, the model was found to provide superior data [20].

### Effectivity and disease progression

DNA from the gastro-intestinal environment of the study animals was extracted using a commercial kit to determine the effect of the drug treatments on the gastric microbiome composition [68,69]. All DNA from the microbiome were analysed with 16S amplicon sequencing using the Illumina Miseq platform. DNA data was analysed using bioinformatics and community profiling was achieved by using MicrobiomeAnalyst. Treatment with the TB drug regime (positive control) had a significant impact on the microbiome diversity compared to *M.tb* infected mice (negative control group), by decreasing the overall diversity of the microbiome. Members of the phylum Bacteroidota were significantly decreased and Firmicutes significantly increased. The effect of Pheroid^®^-based TB therapy on the microbiome did not differ significantly from standard TB therapy. However, the presence of members of the family Saccharimonadaceae was significantly more abundant in the microbiome of mice recovering from Pheroid^®^-based TB therapy, which is positively correlated with immune responses / status [70,71].

Mice from the group treated with the Pheroid^®^ carrier only (excluding APIs) had a significantly higher abundance of *Lactobacillus,* associated with lipid metabolism, in their microbiome. The use of Pheroid^®^ as an alternative for TB drug delivery does not necessarily contribute to improved health of the microbiome during treatment, but may have the potential to stabilize the gastro-intestinal environment after treatment has ceased. Furthermore, the proliferation of commensal bacteria *Parasutterella* observed in the microbiome of mice after treatment with Pheroid^®^ carriers was discontinued. This indicated normalization of the microbiome after exposure (unpublished results). Although the mice treated in this group did not present with a significant reduction of *M.tb* in the lungs, they survived longer than the infected but untreated animals (negative control), suggesting that the Pheroid^®^ formulation may have an immune support function [72].

Interactions between the intestinal microbiome and lungs have been shown to support the body in its fight against TB [73]. Short chain fatty acids (SCFAs) produced by the intestinal microbiome may be instrumental in the construction of an intestinal barrier and respiratory immunity [74]. One may speculate that the fatty acids contained in the Pheroid^®^ may be broken down in the gut to provide SCFAs. Because of the various aberrations of the immune system in the Kramnik mouse model, this may not be the best model to investigate the effect of the Pheroid^®^ carrier on the microbiome and its associated immune support function.

### Strengths and limitations of the Kramnik model and of this study

A distinctive feature of the Kramnik mouse model is its capacity to develop cavitary lesions following aerosol infection, a phenomenon rarely observed in other mouse models [20] and this may bring additional dimensions to the assessment of the efficacy of treatment: the progressive necrotic lesions may harbour great numbers of extracellular bacilli which may lead to similar cavitating lesions as that described in humans [21], but the aberrations in both the humoral and cellular immune systems of these mice may complicate interpretation of the results. While limitations in macrophage pathology in Kramnik mice are well-documented [2], constraints related to B-cell activity have not been adequately demonstrated [75].

The susceptibility of the Kramnik mouse to *M.tb*, driven by the **SST1** locus, predisposes mice to necrotizing TB. This susceptibility allows the model to replicate the extracellular bacterial burden and progressive pathology seen in severe human TB. However, the model’s utility is constrained by several factors:

**1. High inter-individual variability**: As noted by Driver et al. [20], Irwin et al. [19], and Rosenthal et al. [63], significant variability in lung CFU counts following low-dose aerosol infections reduces the statistical power to detect differences between treatment groups. This study mitigated this issue by using a high infection dose of 34,000 CFUs, resulting in well-established bacterial burdens (baseline CFU: 3.63±0.74 log10; pre-treatment CFU: 6.31±0.81 log10). While effective in standardizing infection, this approach introduces a non-physiological scenario that may not accurately reflect human disease dynamics.

**2. Immune dysregulation and macrophage dysfunction**: The Kramnik model’s immune abnormalities are particularly evident in macrophages, which are essential mediators of both innate and adaptive immune responses. Macrophages in C3HeB/FeJ mice exhibit reduced nitric oxide (NO) production, a critical effector molecule in the control of intracellular pathogens. NO deficiency compromises the macrophages’ ability to eliminate intracellular bacteria, allowing for increased replication of *M.tb* in the necrotic lesions [20,63].

Furthermore, macrophages play a pivotal role in antigen presentation, bridging innate and adaptive immunity by presenting antigens via major histocompatibility complex (MHC) molecules [76,77]. In the Kramnik model, impaired macrophage function results specifically in suboptimal antigen presentation via TH2, leading to a dampened humoral immune response. This dysfunction disrupts the activation of B-cells and the subsequent production of antibodies, further compromising the host’s ability to mount an effective immune defense.

**3. High mortality rates**: Consistent with previous studies, this study observed significant mortality in the untreated control group, necessitating euthanasia of several animals ahead of schedule due to *moribund* conditions. This aligns with findings by Lanoix et al. [55], where 29.6% of Kramnik mice succumbed to infection compared to 3.6% of BALB/c mice (p < 0.01). Such high mortality rates reflect the model’s susceptibility, but also limit its applicability to less severe or latent forms of TB.

**4. Challenges in backcrossing**: Stabilizing the genetic background of the Kramnik model requires extensive backcrossing over at least 10 generations. This process is labor-intensive and introduces risks of genetic drift, which can subtly alter immune responses, lesion development, and treatment outcomes. These challenges underscore the need for alternative approaches to maintain genetic consistency.

### Role of SP140 in TB susceptibility and macrophage regulation

SP140, a transcriptional regulator involved in chromatin remodeling, has emerged as a critical locus in macrophage-mediated immunity during TB infection. Studies have demonstrated that SP140 plays a dual role in repressing inappropriate type I interferon responses and promoting the expression of genes critical for pathogen control [29]. Dysregulation of type I interferons has been linked to increased susceptibility to bacterial infections, as excessive interferon signaling can suppress essential pro-inflammatory pathways [29].

SP140-deficient macrophages exhibit reduced transcriptional repression of type I interferon pathways, leading to dysregulated immune responses. This deficiency is compounded by altered chromatin states that impair the activation of key genes involved in macrophage effector functions, such as NO production and phagolysosomal maturation [29]. In the Kramnik model, where macrophage function is already compromised, SP140 plays a crucial role in determining the severity of infection and the host’s ability to control bacterial replication.

The findings by Ji et al. [29] further highlight the translational relevance of SP140. Variants in the SP140 gene have been associated with differential susceptibility to TB in human populations, suggesting that targeting SP140-mediated pathways could offer new therapeutic avenues for enhancing macrophage function and bacterial clearance.

Dysfunction of macrophages occurs in various diseases and often determines the extent of progression, as observed with COVID 2019 and atherosclerosis [78]. Depending on the condition(s) and environmental cues, the functional plasticity of macrophages determine whether these cells exert either a protective or pathogenic effect, which in turn is regulated *via* signal transduction such as Janus kinase–signal transducer and activator of transcription, Wnt and Notch pathways [79]. The pathogenesis of diseases, including autoimmune, neurodegenerative, metabolic, infectious diseases, and cancer is strongly associated with macrophage function and macrophage-targeted therapy has been explored in clinical applications for these diseases [80–82].

The dysfunction of macrophages in the Kramnik model not only exacerbates bacterial replication but also has downstream effects on adaptive immunity. Impaired antigen presentation disrupts the activation of CD4^+^ T-cells, which is critical for coordinating the humoral response. The subsequent lack of robust B-cell activation leads to insufficient antibody production, limiting the host’s ability to neutralize extracellular bacteria. This multifaceted dysfunction underscores the central role of macrophages in orchestrating immune responses and highlights their potential as therapeutic targets.

### Sequencing challenges and genetic insights

The large and repetitive SST1 locus remains a significant challenge for sequencing and genetic analysis. Advances in long-read sequencing technologies, such as those provided by Ji et al. [29], offer potential solutions but require further optimization. In contrast, shorter loci like SP140 and SP110 are more amenable to genetic manipulation and provide clearer insights into immune regulation. SP110, like SP140, is involved in macrophage function and has been shown to regulate bacterial control *via* nuclear signaling pathways [26]. By focusing on these loci, the technical limitations of SST1 can be bypassed and targeted strategies to enhance the translational relevance of the Kramnik model can be developed.

## Conclusion

The Kramnik mouse model (C3HeB/FeJ) has been instrumental in advancing our understanding of TB pathology and treatment. Its unique ability to replicate human-like necrotic granulomas and cavitary lesions provides invaluable insights into advanced disease stages, which are difficult to study in other murine models [21]. However, as highlighted in this study, the model’s strengths are accompanied by significant limitations that complicate its use in evaluating new therapeutic approaches, including carrier systems like Pheroid^®^. These challenges are not only inherent to the Kramnik model, but also reflect broader issues in the use of animal models for translational research.

This study underscores the complexity of evaluating anti-TB therapies using the Kramnik model (C3HeB/FeJ), highlighting both its strengths and limitations. The challenges presented by macrophage dysfunction, impaired NO production, and suboptimal antigen presentation complicate the interpretation of therapeutic outcomes. However, the integration of insights from loci such as SP140 and SP110 offers opportunities to refine the model and enhance its utility in translational research.

SP140, in particular, represents a critical regulator of macrophage function and immune homeostasis, with implications for both TB susceptibility and therapeutic intervention. By addressing the challenges associated with macrophage dysfunction and integrating genetic insights, the potential of the Kramnik model to advance TB research and improve patient outcomes may be harnessed better.

## Acknowledgements

The authors would like to thank Prof. Faans Steyn (in memoriam) at the Statistical Consultation Service Department of the North-West University (NWU) for assisting with statistical data analysis, and Dr. Clarissa Willers for assistance with the manuscript preparation and submission.

## Author contributions

**Conceptualization:** Prof. Anne Grobler, Dr. Theunis Cloete.

**Data curation:** Prof. Anne Grobler, Prof. Markus Depfenhart, Dr. Theunis Cloete, Prof. Yolandy Lemmer.

**Formal analysis:** Prof. Markus Depfenhart, Ms. Drieke van der Merwe, Prof. Anne Grobler.

**Methodology:** Dr. Theunis Cloete, Prof. Anne Grobler, Mr. Kobus Venter.

**Project administration:** Ms. Drieke van der Merwe, Prof. Anne Grobler.

**Resources:** Prof. Anne Grobler (Research budget).

**Statistical analysis:** Prof. Anne Grobler.

**Software:** GraphPad Prism Versions 9 & 10, SPSS.

**Supervision:** Dr. Theunis Cloete, Prof. Anne Grobler.

**Validation:** Prof. Markus Depfenhart.

**Visualization:** Prof. Markus Depfenhart, Prof. Anne Grobler.

**Writing – review & editing:** Prof. Markus Depfenhart, Prof. Anne Grobler, Dr. Theunis Cloete, Prof. Suraj Parihar.

**Writing – original draft:** Prof. Markus Depfenhart, Prof. Anne Grobler, Ms. Drieke van der Merwe.

